# Experience-dependent effects of muscimol-induced hippocampal excitation on mnemonic discrimination

**DOI:** 10.1101/454744

**Authors:** Sarah A. Johnson, Sean M. Turner, Katelyn N. Lubke, Tara L. Cooper, Kaeli E. Fertal, Jennifer L. Bizon, Andrew P. Maurer, Sara N. Burke

## Abstract

Memory requires similar episodes with overlapping features to be represented distinctly, a process that is disrupted in many clinical conditions as well as normal aging. Data from humans have linked this ability to activity in hippocampal CA3 and dentate gyrus (DG). While animal models have shown the perirhinal cortex is critical for disambiguating similar stimuli, hippocampal activity has not been causally linked to discrimination abilities. The goal of the current study was to determine how disrupting CA3/DG activity would impact performance on a rodent mnemonic discrimination task. Rats were surgically implanted with bilateral guide cannulae targeting dorsal CA3/DG. In Exp.1, the effect of intra-hippocampal muscimol on target-lure discrimination was assessed within subjects in randomized blocks. Muscimol initially impaired discrimination across all levels of target-lure similarity, but performance improved on subsequent test blocks irrespective of stimulus similarity and infusion condition. To clarify these results, Exp.2 examined whether prior experience with objects influenced the effect of muscimol on target-lure discrimination. Rats that received vehicle infusions in a first test block, followed by muscimol in a second block, did not show discrimination impairments for target-lure pairs of any similarity. In contrast, rats that received muscimol infusions in the first test block were impaired across all levels of target-lure similarity. Sustained effects of muscimol in disrupting behavioral performance after repeated infusions were verified in a spatial alternation task. At the conclusion of behavioral experiments, fluorescence *in situ* hybridization for the immediate-early genes *Arc* and *Homer1a* was used to determine the proportion of neurons active following muscimol infusion. Contrary to expectations, muscimol increased neural activity in DG. An additional experiment was carried out to quantify neural activity in naïve rats that received an intra-hippocampal infusion of vehicle or muscimol. Results confirmed that muscimol led to DG excitation, likely through its actions on interneuron populations in hilar and molecular layers of DG and consequent disinhibition of principal cells. Taken together, our results suggest disruption of coordinated neural activity across the hippocampus impairs mnemonic discrimination when lure stimuli are novel.

## INTRODUCTION

The neurobiological basis of how distinct events with overlapping features are disambiguated remains a central question in neuroscience. Theory and computational models have often attributed this function to the dentate gyrus (DG) of the hippocampus, based on its densely packed granule cell population in which a small number of afferent neurons project onto many more granule cells (Amaral et al., 1990, 2007; Patton and McNaughton, 1995; Amaral and Lavenex, 2007; Treves et al., 2008), as well as its relatively sparse neural activity (McNaughton and Morris, 1987; Treves and Rolls, 1992; O’Reilly and McClelland, 1994; Rolls and Kesner, 2006; Leutgeb and Leutgeb, 2007; Treves et al., 2008; Schmidt et al., 2012; Santoro, 2013; Kesner and Rolls, 2015; Knierim and Neunuebel, 2016).

Hippocampal contributions to the orthogonalization of similar inputs have been assessed with tasks that require discrimination of a previously viewed target image from lure images, which range in similarity to the target. These paradigms are now commonly referred to as ‘mnemonic similarity’ tasks (Stark et al., 2013, 2015; Huffman and Stark, 2017; Stark and Stark, 2017). Functional neuroimaging studies using these tasks support earlier predictions that the ability to accurately resolve a target from similar lures is linked to activation in the DG and CA3 sub-regions of the hippocampus (Kirwan and Stark, 2007; Bakker et al., 2008; Lacy et al., 2011 p.2; Kirwan et al., 2012; Motley and Kirwan, 2012; Reagh and Yassa, 2014; Reagh et al., 2017). Furthermore, mnemonic discrimination deficits in older adults and individuals with amnestic mild cognitive impairment (aMCI) correlate with altered activation of CA3/DG (Yassa et al., 2010, 2011a, 2011b; Bakker et al., 2012, 2015; Reagh et al., 2018), decreased integrity of perforant path fiber tracts, which provide direct inputs from parahippocampal cortical structures to DG (Bennett et al., 2015; Bennett and Stark, 2016), and changes in CA3/DG volume (Doxey and Kirwan, 2015).

We have recently validated a rodent version of the mnemonic similarity task (Johnson et al., 2017), in which elevated activity in CA3 is linked to impaired performance (Maurer et al., 2017). Discrimination performance in this continuous, forced-choice paradigm is contingent on the proportion of visible features shared between the learned target object and lure objects. Additionally, aged rats are impaired in distinguishing the target from similar lures, but not distinct lures, which directly parallels previous findings from older adults (Toner et al., 2009; Yassa et al., 2011a, 2011b; Ryan et al., 2012; Holden et al., 2013; Stark et al., 2013, 2015; Pidgeon and Morcom, 2014; Reagh et al., 2016, 2018; Huffman and Stark, 2017; Stark and Stark, 2017; Trelle et al., 2017).

Based on models of hippocampal network function (O’Reilly and McClelland, 1994; Yassa and Stark, 2011; Kesner and Rolls, 2015; Knierim and Neunuebel, 2016; Leal and Yassa, 2018), and data from neuroimaging studies mentioned above, we hypothesized that disrupting neural activity in CA3/DG would selectively impair discrimination of a learned target object from similar lure objects, when the target and lures share a relatively high percent feature overlap (i.e. more than 70%; Johnson et al., 2017). We tested this prediction in the current experiments by infusing the GABAA agonist muscimol through guide cannulae targeting the dorsal CA3/DG of young adult rats prior to mnemonic discrimination testing. At the completion of behavioral testing, the effects of muscimol in the hippocampus were examined by labeling mRNA transcripts of the activity dependent immediate-early genes *Arc* and *Homer1a*. A previous study showed that muscimol infusions into the medial prefrontal and perirhinal cortices blocks expression of *Arc*, verifying the efficacy of neural inactivation through GABAA agonism (Hernandez et al., 2017). Surprisingly, in the current study, muscimol infusions centered on dorsal CA3/DG did not block immediate-early gene expression, but rather led to *increased* activity within the DG. This likely occurred through inactivation of adjacent interneuron populations in the hilus and DG molecular layers, which in turn disinhibited DG granule cells. Thus, a critical conclusion of these studies is that it is essential to confirm the effects of pharmacological agents or other functional manipulations using *both* behavioral and neural read-outs (Allen et al., 2015; Smith et al., 2016). Importantly, this hyperactivity within DG, which recapitulates sub-convulsive seizures observed in Alzheimer’s disease (Palop and Mucke, 2009, 2010; Vossel et al., 2013, 2016), impaired mnemonic discrimination performance across all target-lure pairs irrespective of their similarity. Additional testing revealed that muscimol infusions impaired mnemonic discrimination performance only when lure objects were novel.

## MATERIALS AND METHODS

### Animals

A total of 38 young adult male Fischer 344 x Brown Norway F1 hybrid rats (NIA colony, Taconic; 4-6 months of age at arrival) were used as subjects (Exp.1: 10 rats; Exp.2: 20 rats, 3 excluded due to blocked or misplaced cannulae; Verification of neural effects of muscimol: 8 rats, 2 excluded due to misplaced cannulae). Rats were single-housed in standard Plexiglas cages and maintained on a 12-h reverse light/dark cycle (lights off 8:00 am). All manipulations took place in the dark phase, 5-7 days per week at approximately the same time each day. Rats received 20±5 g moist chow (~39 kcal; Teklad LM-485, Harlan Labs) daily and drinking water *ad lib*. Shaping began once they reached 85% of their initial body weights on restricted feeding, which provided appetitive motivation in behavioral tasks. All procedures were carried out in accordance with the NIH *Guide for the Care and Use of Laboratory Animals* and approved by the Institutional Animal Care and Use Committee at the University of Florida.

### Verification of cannulae placements and neural effects of muscimol

After completing infusions and mnemonic discrimination testing in Experiments 1 and 2 (Fig.1A), brain tissue was collected for histological verification of cannulae placements. Effects of muscimol on neural activity within the diffusion radius of the drug were also assessed by visualizing mRNA of the immediate-early gene *Arc* with fluorescence *in situ* hybridization (FISH). All rats from Exps.1 and 2 received intra-hippocampal muscimol infusions (1μL, 1mg/mL; see subsequent sections for details) and were returned to their home cages for 30 min. Rats performed a 10-min mnemonic discrimination test, were transferred to a glass bell jar for deep anesthesia with isoflurane (Isothesia, Henry Schein), and were sacrificed by rapid decapitation. Brains were extracted and snap-frozen in chilled isopentane (Acros Organics, Fisher Scientific, Pittsburgh, PA). Tissue blocks containing 2 to 4 brains each were sectioned (20 μm) on a cryostat (Microm HM550; ThermoFisher Scientific, Waltham, MA), thaw-mounted on Superfrost Plus slides (Fisher Scientific), and stored at −80°C in sealed slide boxes. Dorsal hippocampal sections were DAPI-stained (ThermoFisher Scientific) and imaged with fluorescence microscopy (Keyence; Itasca, IL) to map position of guide cannulae (Fig.1B-D). Slides with visible cannulae tracks were then processed with FISH to verify effects of muscimol on neural activity at the infusion site.

**Figure 1.**
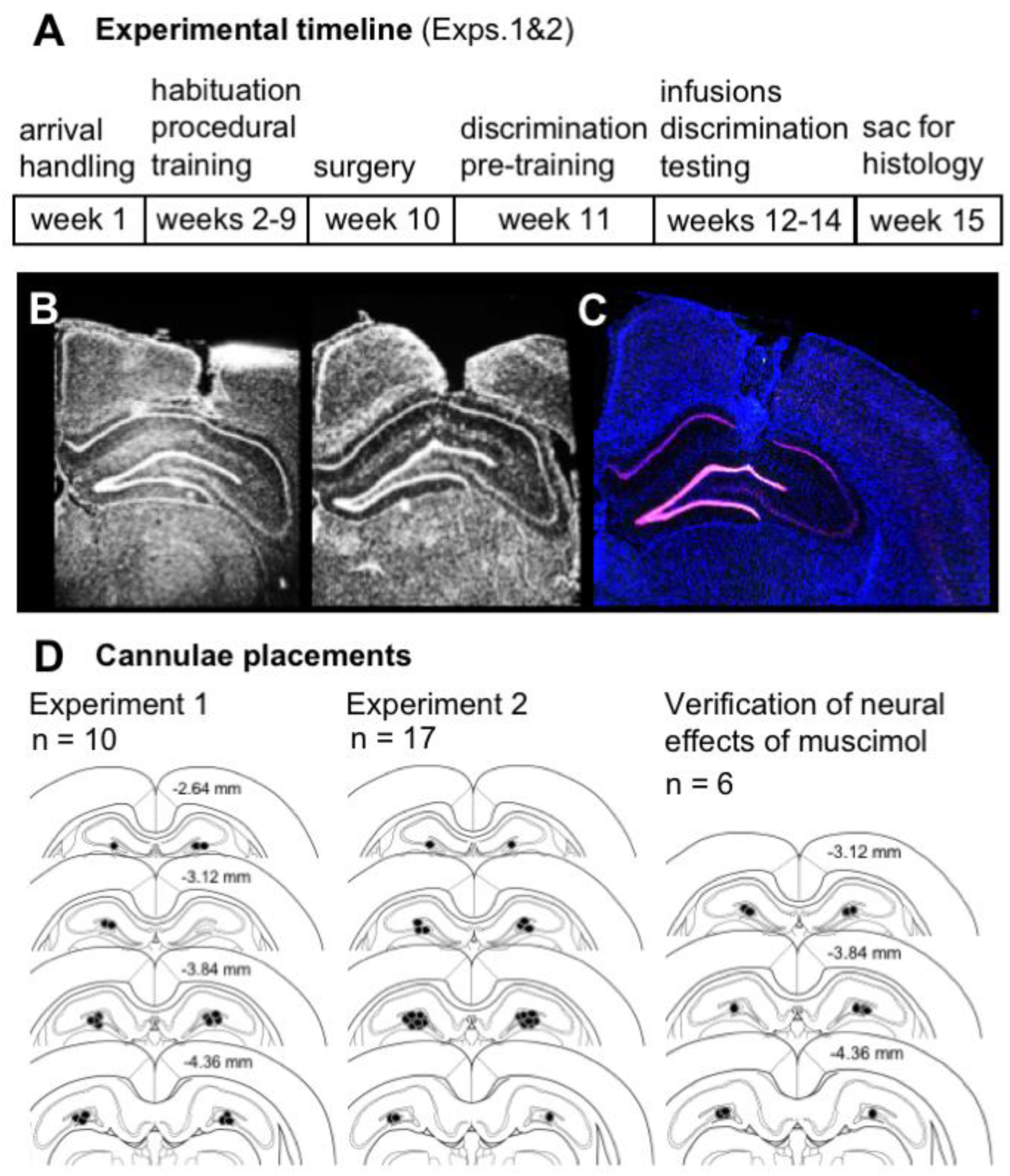
(**A**) Timeline of experimental manipulations and histological assessment of dorsal CA3/DG cannulae placements from Experiments 1 and 2. (**B**) Representative images show DAPI-stained sections (converted to grayscale) with tracts of 22G stainless steel guide cannulae positioned above the dorsal CA3/DG at target coordinates, relative to Bregma: AP −4mm, ML ±3mm, DV −2.6mm from skull surface. While no visible damage was observed in most rats, 28G microinjectors protruded an additional 1mm below cannulae tips, centering the infusion between DG upper and lower blades. (**C**) Representative image from Exp.1 brain section processed with fluorescence *in situ* hybridization (FISH) to label *Arc* mRNA (Cy3; red channel), counter-stained with DAPI. Strong induction of the immediate-early gene *Arc* is evident in both upper and lower blades of DG. (**D**) Schematics show position of foci of infusions based on cannulae tracts in each rat from Exp.1 (n=10), Exp.2 (n=17), and separate study carried out to verify effects of vehicle vs. muscimol infusion on Arc mRNA expression as a read-out of neural activity (n=6). Numerical values on brain sections indicate AP coordinate, relative to Bregma.

FISH for *Arc* mRNA was carried out as previously described (Guzowski et al., 1999; Guzowski and Worley, 2001; Burke et al., 2012; Hernandez et al., 2017, 2018; Maurer et al., 2017). Briefly, digoxigenin (DIG)-labeled riboprobes were generated with a commercial transcription kit (Riboprobe System, P1440; Promega, San Luis Obispo, CA) and DIG RNA labeling mix (Roche REF# 11277073910; MilliporeSigma, St. Louis, MO) from a plasmid template containing full-length 3.0 kb *Arc* cDNA under the T7 promoter (generously provided by Dr. A. Vazdarjanova; Augusta University, Augusta, GA). Slides were hybridized overnight, then incubated with anti-Digoxigenin-POD (Roche REF# 11207733910, MilliporeSigma) overnight. *Arc*-labeled transcripts were visualized with Cy3 (TSA Cyanine 3 System, NEL744A001KT; PerkinElmer, Waltham, MA) and sections were counter-stained with DAPI. Stitched low magnification flyover images were collected by fluorescence microscopy (Keyence) with a 2X objective. Initial inspection of *Arc* mRNA labeling in tissue from Exp.1 and 2 rats revealed strong induction of neuronal activity in the DG (Fig.1C).

An additional experiment was therefore designed to explicitly test the effect of muscimol infusions in dorsal CA3/DG on neuronal activity in surrounding hippocampal sub-regions. Rats (n=8) were placed on a restricted feeding schedule to match conditions of prior experiments. After reaching 85% of their initial body weights (~2 weeks), rats were surgically implanted with bilateral guide cannulae targeting dorsal CA3/DG (see subsequent sections for details) and allowed 1 week post-op recovery. Throughout this period from arrival to complete recovery, rats were handled extensively by experimenters during daily weighing and feeding to match conditions of prior experiments. Rats were acclimated to infusion procedures for 3 consecutive days (days 1-3). This acclimation included transport to the procedure room, gentle restraint and handling by experimenters during the infusion, removal and cleaning of dummy stylets, and insertion of microinjectors. On day 4, all rats received a vehicle infusion to provide habituation to the infusion procedure. Handling continued for an additional 2 days (days 5-6), then on day 7 rats were randomized to receive either vehicle or muscimol. Infusions were given as for previous experiments, and rats were returned to their home cage for 40 min prior to collection of brain tissue. Blocks containing brains from each infusion condition were processed as in previous experiments. To provide a read-out of baseline neural activity in addition to activity induced by muscimol infusions, a dual-label FISH protocol was used to visualize both *Homer1a* and *Arc* mRNAs. The temporal dynamics of *Homer1a* transcription following neuronal activity are offset from those of *Arc*, which allows identification of cells active approximately 60 min (*Homer1a* cytoplasm labeling) versus 30 min (*Arc* cytoplasm labeling + *Homer1a* foci labeling) prior to tissue collection (Vazdarjanova et al., 2002; Marrone et al., 2008). Probes were generated from plasmids containing the full-length *Homer1a* cDNA under the T7 promoter (provided by Dr. A. Vazdarjanova; Augusta University, Augusta, GA) and *Arc* cDNA (as for Exps.1 and 2), then visualized with Cy3 (*Homer1a*; as for Exps.1 and 2, PerkinElmer) and fluorescein (*Arc*; TSA Fluorescein System, NEL701A001KT; PerkinElmer). Stitched flyover images of the dorsal hippocampus and z-stacks from regions of interest were collected by fluorescence microscopy (Keyence) with 2X and 40X objectives, respectively. Cannulae placements were verified from flyover images (Fig.1D; 2 vehicle rats excluded due to cannulae posterior to desired target). Neurons of CA1 and CA3 showing immediate-early gene localization indicative of baseline activity (*Homer1a* cytoplasm) versus infusion-induced activity (*Arc* cytoplasm + *Homer1a* nuclear foci) were counted manually using ImageJ software and a custom plug-in. Given clear patterns of expression observed on first pass, and challenges posed to manual counting by the densely packed granule cell layer, DG mRNA expression was quantified with densitometry, also using ImageJ.

### Habituation and procedural training

In Exps.1 and 2, object discrimination tasks were carried out in an L-shaped maze bounded by a start area and choice platform, as previously described (Fig.2A; Johnson et al., 2017; Maurer et al., 2017). Procedures for shaping and object discrimination training were identical to those described previously (Johnson et al., 2017). Briefly, rats were habituated to the maze over the course of 1-3 days by free-foraging for scattered pieces of Froot Loop cereal (Kellogg’s; Battle Creek, MI), which served as the food reward throughout experiments. Rats were next shaped to alternate between the start area and choice platform by providing food reward in each location. After reaching a criterion of 32 alternations within 20 min, rats were trained on procedural aspects of the forced-choice object discrimination task with a pair of ‘standard’ unrelated objects, followed by two pairs of LEGO objects (Fig.2B). Objects of one LEGO pair were perceptually distinct, sharing 38% front-facing visible features, while the other pair were perceptually similar, sharing 63% front-facing features. Detailed descriptions of these stimuli and calculations of their feature overlap are provided in Johnson et al. (2017). For each pair, rats learned to identify one object as the target (S+), placed over the food well baited with food reward, while the alternate object was a ‘lure’ (S-), placed over the empty food well. Training proceeded in daily sessions of 32 trials. In each trial, rats exited the start area, traversed the maze to the choice platform, and opted to displace one of the two objects covering the food wells. If the target object (S+) was selected, rats consumed the revealed food reward and returned to the start area to receive a second food reward. If the lure object (S-) was selected, both objects and the food reward were quickly removed from the choice platform and the rat did not receive a food reward in the start area. The side of the baited food well on the choice platform was pseudo-randomized across trials to provide an equal number of left- and right-rewarded trials in each session. Further, the object serving as the target for each pair and the order in which LEGO pair training took place (distinct vs. similar) were counterbalanced across rats. Training on each pair was considered complete when rats reached a criterion of ≥26 correct responses out of 32 trials (≥81.3%) on a single training session. A subset of rats from Exp.1 completed this training as part of a prior study (n=5, Johnson et al., 2017).

**Figure 2.**
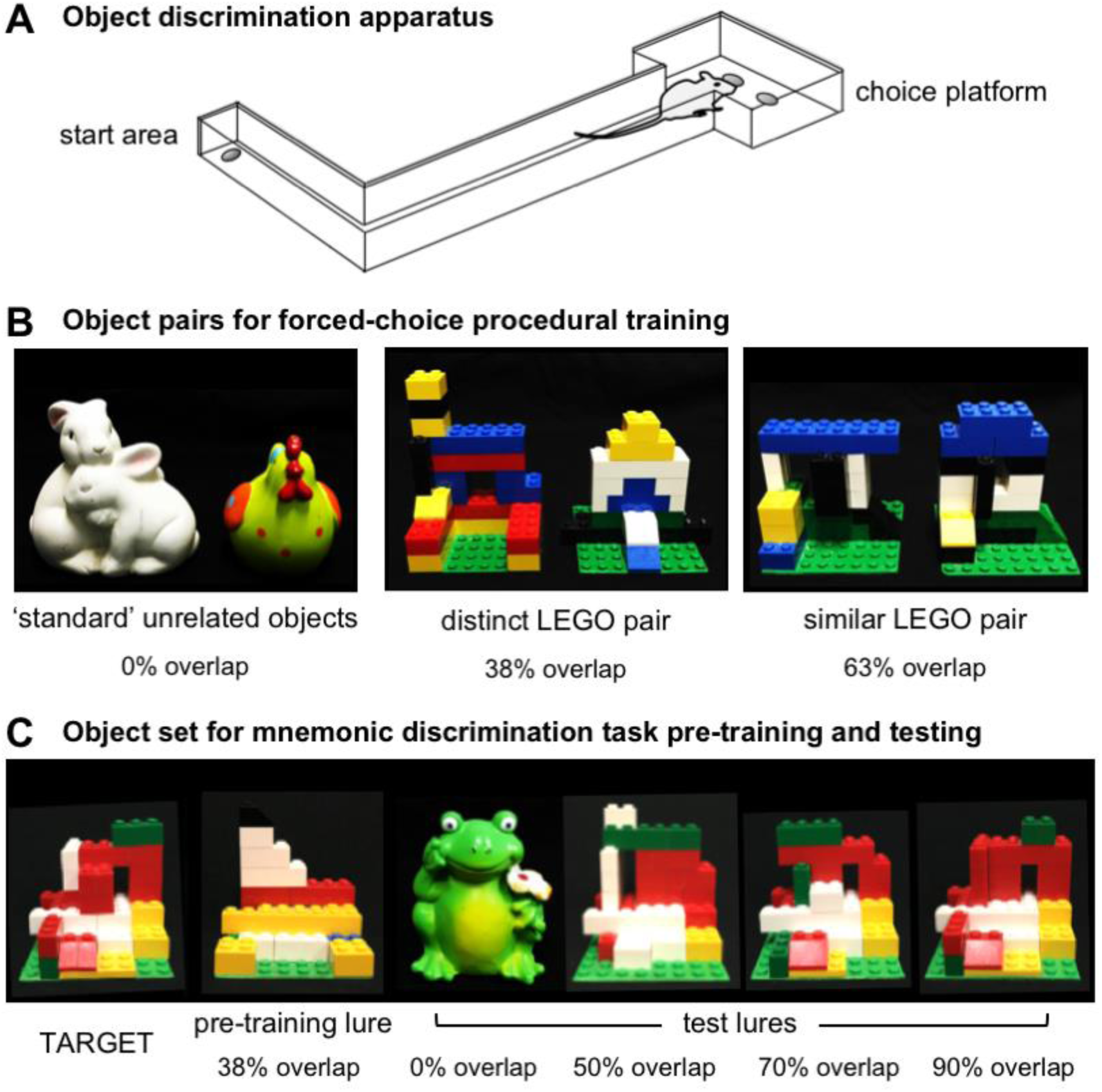
Apparatus and object stimuli used in procedural training and the rodent version of the mnemonic similarity task, as previously described by Johnson et al. (2017) (**A**) Object discrimination training and testing were carried out in an L-shaped maze. Food rewards were hidden in recessed wells in the choice platform, covered by object stimuli. Rats alternated between the start area and choice platform on each discrimination trial. (**B**) Object pairs used in procedural training. All rats were first trained to criterion of >81.3% (>26/32 correct trials) in distinguishing a pair of standard unrelated objects that shared no feature overlap, then were trained to the same criterion on a distinct pair of LEGO objects that shared 38% visible front-facing features, and a similar pair of LEGO objects that shared 63% visible front-facing features. The object of each pair serving as the target (S+) and the order of training on distinct vs. similar LEGO pairs was counter-balanced across rats in each experiment. (**C**) Objects used in the rodent version of the mnemonic similarity task. Left side of panel shows target (S+) object and LEGO pre-training lure object (S-) that shared 38% front-facing visible features. Rats were trained to discriminate this target from the pre-training lure to a criterion of >81.3% (>26/32 correct trials) before moving on to mnemonic discrimination tests. Right side of panel shows each of 4 LEGO test lure objects in order of increasing similarity, or percent visible feature overlap: a distinct, standard lure object that shared 0% overlap with the target, and LEGO lure objects that shared 50, 70, and 90% overlap with the target.

### Surgery

After completing procedural training, rats were surgically implanted with bilateral guide cannulae in the dorsal hippocampus, targeting the pyramidal cell layer of proximal CA3 and the surrounding DG. While maintained on 1-3% isoflurane anesthesia (Isothesia, Henry Schein, Dublin, OH), a longitudinal incision was made to expose and clean the skull surface. Four stainless steel bone screws were placed (flat point, #000-120, 1/8”; Antrin Miniature Specialties) and stainless steel guide cannulae (22 gauge; Plastics One, Roanoke, VA) were positioned at, relative to Bregma, AP −4mm, ML ±3mm, DV −2.6mm from skull surface. The dorsal-ventral positions of cannulae tips were 1mm above the ultimate infusion sites, as microinjectors protruded 1mm below this coordinate. Cannulae were secured to the skull surface and anchor screws with dental cement (Teets, Patterson Dental, Tampa, FL). Dummy stylets (Plastics One) were secured to prevent contamination and maintain patency. Meloxicam (1-2mg/kg s.c.; Metacam, Boehringer Ingelheim, Vetmedica Inc., St. Joseph, MO) was given for pre- and post-op analgesia. Rats were allowed a recovery period of 1 week prior to resuming behavioral experiments.

### Intra-hippocampal infusions

Infusions of the GABAA receptor agonist muscimol (1mg/mL; Sigma-Aldrich, St. Louis, MO) or vehicle (0.9% sterile physiological saline) were given in a 1μL volume at a rate of 0.25μL/min. Internal stainless steel microinjectors (28 gauge, Plastics One) protruding 1mm below implanted guides were connected via polyethylene tubing (PE50, Plastics One) to 10μL syringes (Hamilton, Franklin, MA) mounted in a microinfusion pump (Harvard Apparatus, Holliston, MA). Tubing was backfilled with autoclaved water. A 1-μL bubble was aspirated to create a barrier between backfill and infusate, and permit confirmation by visual inspection that the correct volume of infusate had been delivered. Microinjectors were left in place for 2 min after the infusion to allow dispersion of the drug. Rats were then returned to the home cage for 30 min prior to beginning behavioral testing.

### Mnemonic discrimination task

After their 1-week recovery period, rats were trained to identify a new LEGO object that would serve as the target (S+) throughout mnemonic discrimination testing (Fig.2C). Pre-training was carried out with this target and a perceptually distinct LEGO lure object (S-; Fig.2C). This pair shared 38% front-facing features, comparable to the distinct LEGO pair used in procedural training. After reaching criterion of ≥81.3% correct responses on a single session, rats were given 2 days off before their first mnemonic discrimination test. Tests then proceeded every 3 days (day 1 test, days 2-3 off) to provide a drug wash-out period between infusions (Bañuelos et al., 2014; McQuail et al., 2016).

Experiment 1 was designed to determine if intra-hippocampal muscimol infusions selectively impaired rats’ abilities to discriminate a known target object from similar lure objects. Infusions and tests proceeded over 3 blocks. Each rat completed one test with vehicle and one test with muscimol in each block, with order of infusion conditions pseudo-randomized across subjects in a Latin square design (Fig.5A). As results of Exp.1 revealed muscimol differentially influenced performance across test blocks, Exp.2 was designed to probe this interaction. Infusions and tests were given in 2 blocks, with 3 tests per block. In the first block, separate groups of rats received infusions of vehicle or muscimol on all 3 tests. In the second block, rats received the reverse infusion condition on all 3 tests (Fig.6A).

Mnemonic discrimination tests were carried out as previously described (Johnson et al., 2017). Each session comprised 50 trials: 10 with an entirely distinct standard lure object (frog figurine, 0% feature overlap), 10 with each of 3 perceptually similar LEGO lure objects, sharing 50, 70, and 90% front-facing features, respectively, and 10 with an identical copy of the target object (Fig.2C). Trials with the identical target were included as a control condition, to verify rats were not using olfactory cues from the maze, objects, or food rewards to guide response selection. Test sessions were recorded with a webcam mounted above the choice platform. Response selection behavior and reaction times were then scored offline with custom software (Collector/Minion; Burke/Maurer Labs, Gainesville, FL).

In Exp.2, a behavioral control task was included following mnemonic discrimination tests to assess patency of guide cannulae and continued effectiveness of muscimol in altering behavior after repeated infusions. Rats were trained to spatially alternate on a figure-8 maze to a criterion of 10 consecutive correct alternations, then were infused immediately with 1μL muscimol (1mg/mL) and re-tested for spatial alternation abilities after 30 min. Three days later, rats were returned to the maze, pre-trained to the same criterion, immediately infused with 1μL vehicle, and tested after 30 min. Infusions were given in this sequence for all rats as spatial alternation abilities are most reliant on neural activity in the dorsal hippocampus prior to over-training and automation of responding (Stevens and Cowey, 1973; Rawlins and Olton, 1982).

### Statistical analyses

Analyses were performed with the Statistical Package for the Social Sciences (SPSS) v25 for Windows. Immediate-early gene data were analyzed with repeated measures ANOVA, with gene, region, and time point as within subjects factors, and infusion condition as a between subjects factor. Behavioral data were also analyzed with repeated measures ANOVA, with test block, test day, and lure object as within subjects factors. Infusion condition was assessed within subjects in Exp.1, and between subjects in Exp.2. Significant effects and interactions were followed with simple contrasts. When relevant, performance was compared to chance levels (i.e., 50% correct responses) with one-sample t-tests. Choice of statistical test was dictated by assumptions of normality, assessed with Shapiro-Wilk tests, and homogeneity of variances, assessed with Levene’s tests. P-values < 0.05 for ANOVA or Bonferroni-corrected based on the number of comparisons performed were considered statistically significant.

## RESULTS

### Muscimol increased neural activity in DG

After noting in Exps.1 and 2 that muscimol appeared to induce, rather than silence, DG neural activity, a study verifying expression of activity-dependent immediate-early genes *Arc* and *Homer 1a* was carried out to clarify the effect of GABAA agonism in dorsal CA3/DG on neural activity across hippocampal subregions. For analyses, image stacks were captured at high magnification from CA1 and CA3 near the site of cannulae implantation (Fig.3A), and stitched flyover images were captured at low magnification of the entire DG (Fig.3B). Representative z-stacks that reflect the distribution of *Arc* and *Homer1a* signal after vehicle versus muscimol infusion are shown in Fig.3C-E. For CA1 and CA3, all DAPI-labeled neuronal nuclei present in the median 20% of z-stacks were counted, then classified based on sub-cellular distribution of mRNA as active at baseline or activated by infusion. Percentages of active neurons at each of these time points are shown in Fig.3F. For DG, circular cursors spanning the width of the granule cell layer were placed along upper and lower blades (Fig.1B) and mean Integrated density values were averaged across each blade (Fig.3G).

Inspection of DG images confirmed muscimol, but not vehicle, generated excitation of granule cells (Fig.3B/E). While immediate-early gene expression was also evident in CA1 and CA3 principal cells (Fig.3C/D), neural activity in these sub-regions did not differ based on the infusate delivered. Repeated measures ANOVA with hippocampal subregion and time point entered as within subjects factors and infusion condition as a between subjects factor revealed a statistically significant effect of time point on proportion of neurons active (*F*_(1,4)_ = 24.3, *p* < 0.008). Specifically, receiving an intra-hippocampal infusion of vehicle or muscimol increased the percent of neurons active in both CA1 and CA3 relative to baseline (Fig.3F, no main effect of infusion condition: *F*_(1,4)_ = 0.39, *p* = 0.565). All other main effects and interactions were not statistically significant (*p*’s > 0.19). For DG, initial comparison of *Arc* and *Homer1a* signals sampled on green and red channels showed no difference between levels of the two mRNA transcripts (no main effect of IEG: *F*_(1,7)_ = 1.08, *p* = 0.334), therefore only the *Arc* data were analyzed within this subregion (upper vs. lower blade) for an effect of infusion condition. Repeated measures ANOVA revealed a significant main effect of infusion condition (*F*_(1,3)_ = 16.7, *p* < 0.027), indicating that muscimol infusion led to greater *Arc* expression in both upper and lower blades of DG, compared to rats with vehicle infusion (no main effect of subregion or infusion x subregion interaction, *p*’s > 0.328). Together these data indicate that muscimol infusions at the coordinates used in the current study produced DG hyperexcitation, while not altering the portions of active principal neurons in CA3 and CA1. Thus, the results of behavioral experiments must be considered in a framework of aberrant granule cell firing rather than the silencing of hippocampal principal cells.

**Figure 3.**
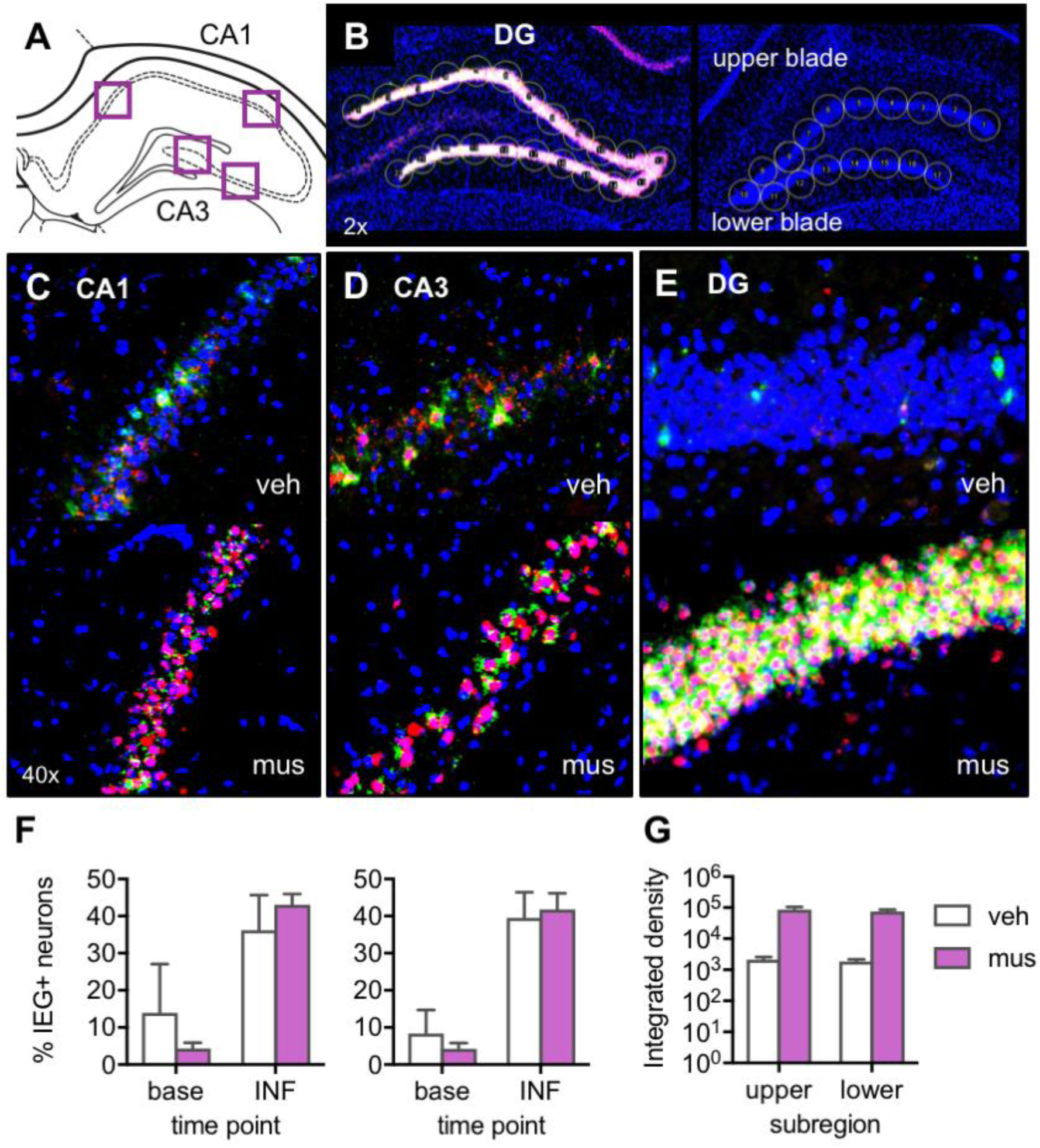
Verification of the effect of muscimol infusions on neural activity across hippocampal subregions with fluorescence *in situ* hybridization (FISH) for the immediate-early genes (lEGs) *Arc* and *Homer 1a.* (**A**) Regions of interest from which z-stacks were captured in CA1 and CA3 for quantification of lEGs by manual counting in ImageJ. (**B**) Representative images from a muscimol-infused rat (left) and vehicle-infused rat (right) show schematic of cursors aligned across upper and lower blades of DG for densitométrie quantification of lEGs using ImageJ software. (**C-E**) Representative z-stacks show *Arc* (fluorescein; green channel) and *Hornería* (Cy3; red channel) labeling from a vehicle (veh)-infused and muscimol (mus)-infused rat; (**C**) CA1, (**D**) CA3, and (**E**) DG. (**F**) Plots show mean ± SEM percent IEG-positive DAPI-stained neuronal nuclei reflecting cells active at baseline (base), 60 min prior to sac (*Hornería* cytosol), versus cells activated by the infusion (INF), 30 min prior to sac (*Hornería* nuclear foci + *Arc* cytosol). Both infusion conditions increased percent neurons active in CA1 and CA3 relative to baseline (main effect time point: *F*_(1,4)_ = 24.3, *p* < 0.008; no main effect infusion condition: *F*_(1,4)_ = 0.39, *p* = 0.565). (**G**) Mean ± SEM integrated density of *Arc* mRNA signal quantified in DG upper and lower blades. Muscimol infusion led to greater *Arc* expression in both DG subregions (main effect infusion condition: *F*_(1,3)_ = 16.7, *p* < 0.027).

### Procedural training

Figure 4 shows the number of incorrect trials completed by rats in each experimental group prior to reaching criterion in procedural training (Fig.4A), and pre-training with the target object used in mnemonic discrimination tests (Fig.4B). Rats from Exp.1 and both infusion groups in Exp.2 did not differ in the amount of training required (*F*_(2,25)_ = 0.79, *p* = 0.46, no group x object pair interaction: *F*_(6,75)_ = 1.24, *p* = 0.30). The main effect of object pair (standard versus LEGO objects), however, significantly affected the number of incorrect trials made prior to criterion (*F*_(3,75)_ = 68.6, *p* < 0.001). Consistent with our prior studies (Johnson et al., 2017), rats required fewer trials to reach criterion on the standard object pair versus all other pairs (simple contrasts, α = 0.05/6 = 0.008, *p*’s < 0.001), but a greater number of trials to reach criterion on the similar LEGO pair versus all other pairs (*p*’s < 0.001). Critically, rats reached criterion performance for the distinct LEGO pair used in initial procedural training, before surgery, and the mnemonic discrimination task pre-training pair, after surgery, in a comparable number of incorrect trials (*F*_(1,29)_ = 0.28, *p* = 0.60). This indicates hippocampal cannulation and post-op recovery did not reduce procedural knowledge of the discrimination task, or rats’ abilities to learn a new target object.

**Figure 4.**
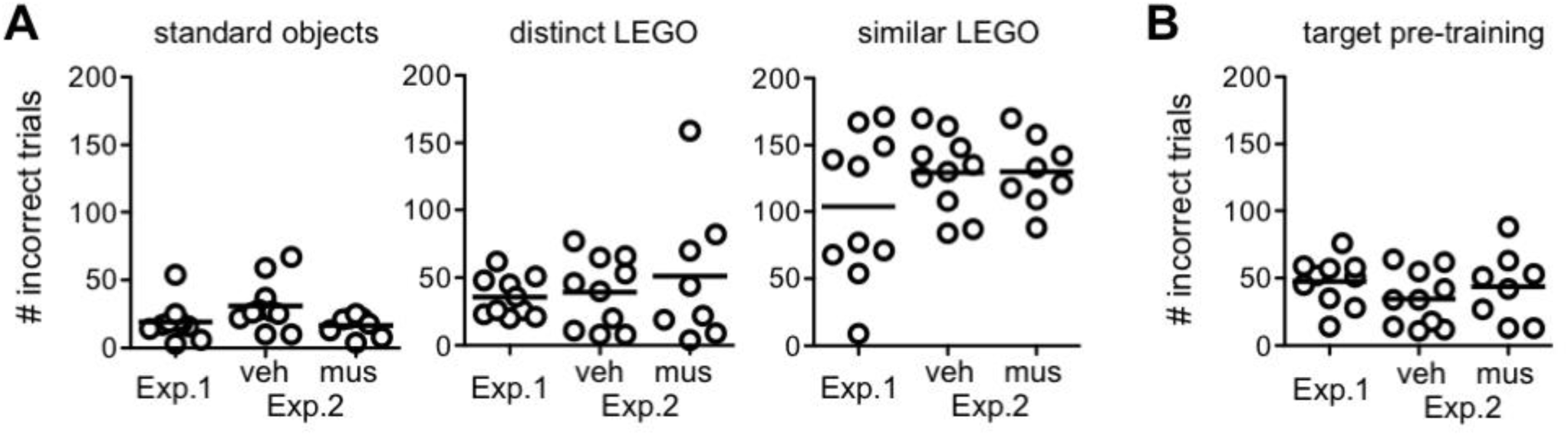
Procedural object discrimination training required to reach a criterion performance level of >81.3% (>26/32 correct responses) on a single daily session in Experiments 1 and 2. (**A**) Number of incorrect responses required to reach criterion on object pairs used for procedural discrimination training in rats from Exp.1, and rats assigned to groups that received vehicle infusions first (veh) or muscimol infusions first (mus) across two test blocks in Exp. 2. Irrespective of experiment or group, all rats learned to accurately identify the target of the standard object pair after fewer incorrect trials, relative to all other pairs (simple contrasts, *p*'s < 0.001). Conversely, all rats required a greater number of incorrect trials to reach criterion on the similar LEGO pair (simple contrasts, *p*'s < 0.001). (**B**) Number of incorrect responses made prior to reaching criterion in identifying the target (S+) object relative to the distinct lure object in pre-training for mnemonic discrimination tests. Amount of training required did not differ by experiment or group. Additionally, amount of training required to reach criterion on this pair, after cannulation surgery, did not differ from that required to reach criterion on the distinct LEGO pair prior to surgery (*p* = 0.60).

### Exp.1: Muscimol impaired discrimination, irrespective of target-lure similarity

Target-lure discrimination tests included a control object pair, consisting of 2 identical target objects, to ensure that rats were not smelling the food reward or using another latent variable to solve discrimination problems. Rats performed at chance levels (i.e., 50% correct) on these control trials. A repeated measures ANOVA comparing performance on these trials indicated no significant difference across infusion conditions (*F*_(2,18)_ = 0.16, *p* = 0.85). Further, one-sample t-tests against a hypothetical mean of 50% showed performance did not differ from chance levels (veh: *t*_(9)_ = 0.51, *p* = 0.62; mus: *t*_(9)_ = −0.05, *p* = 0.96). Control trials were therefore excluded from all subsequent analyses.

Performance on mnemonic discrimination tests for target-lure problems with ≤90% feature overlap is shown in Figure 5. A timeline of Exp.1 infusions and tests given across 3 test blocks is shown in Figure 5A. When performance values were collapsed across test blocks (Fig.5B), a repeated measures ANOVA with infusion condition and lure as within subjects factors revealed a significant main effect of lure (*F*_(3,27)_ = 44.1, *p* < 0.001). Specifically, rats made fewer correct responses on trials with LEGO lures (50-90% overlap), relative to the standard object lure (0% overlap; simple contrasts: *p’s* < 0.01). However, performance across lure objects did not differ based on infusion condition (main effect of infusion: *F*_(1,9)_ = 2.73, *p* = 0.13, lure x infusion interaction: *F*_(3,27_) = 0.38, *p* = 0.77).

**Figure 5.**
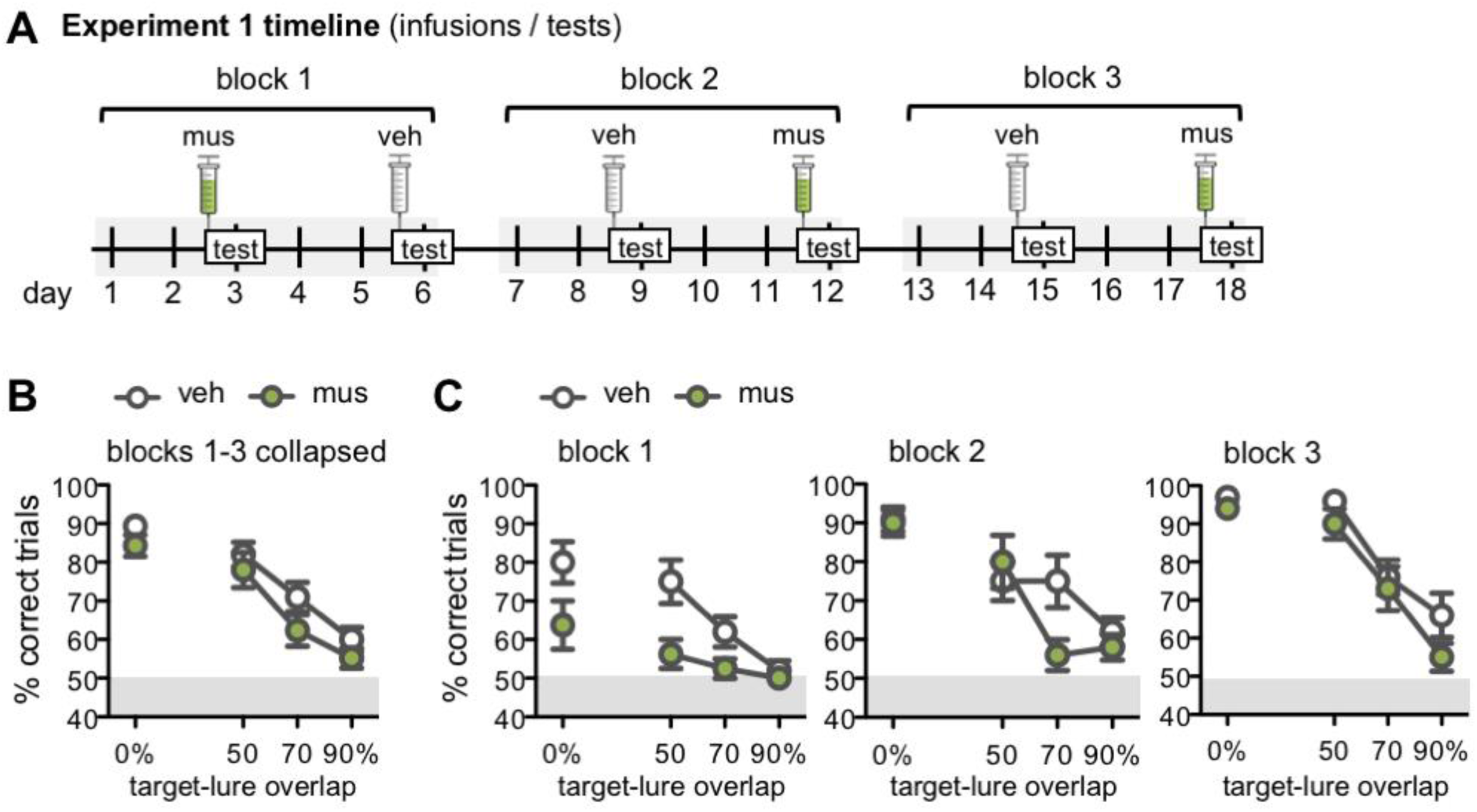
(**A**) Timeline of infusions and mnemonic discrimination tests administered in Experiment 1. All rats (n=10) received a vehicle (veh) and muscimol (mus) infusion in each of 3 tests blocks. Order of veh and mus infusions in each test block was randomized across rats with a Latin square design, therefore this timeline shows one example permutation of infusion order. Tests took place every 3 days, with 2 wash-out days on which rats remained in their home cages and did not complete any behavioral training or testing. Infusions were administered 30 min prior to the beginning of each test. (**B/C**) Performance (% correct trials) on mnemonic discrimination tests in Exp.1 plotted by trial type, with each of the 4 lure objects sharing 0, 50, 70, or 90% visible front-facing features (target-lure overlap). Test performance on days with vehicle infusions (veh) designated by open circles, and on days with muscimol infusions (mus) by filled circles. (**B**) Mean ± SEM performance on mnemonic discrimination tests collapsed across 3 test blocks, to a total of 30 trials with each lure object under each infusion condition. Rats made fewer correct responses on trials with LEGO lures (50-90% target-lure overlap), relative to the standard object lure (0% overlap; main effect lure: *F*_(3,27)_ = 44.1, *p* < 0.001; simple contrasts: *p*’*s* < 0.01). (**C**) Mean ± SEM performance plotted for each of the 3 test blocks (10 trials / lure object / infusion condition). Muscimol impaired discrimination when lure objects were novel, in block 1, across all lures (main effect infusion: *F*_(1,7)_ = 16.7, *p* < 0.005). By block 2, muscimol impaired performance only on the 70% target-lure overlap problem (infusion × lure: *F*_(3,27)_ = 2.95, *p* < 0.05). No difference in performance between infusion conditions was observed in block 3.

Follow-up analyses compared discrimination performance across test blocks to determine if the effect of muscimol varied with increasing task experience (Fig.5C). Repeated measures ANOVA revealed significant main effects of block (*F*_(2,14)_ = 21.4, *p* < 0.001), lure (*F*_(3,21)_ = 37.1, *p* < 0.001), and infusion condition (*F*_(1,7)_ = 10.9, *p* < 0.013), in addition to a significant block x lure interaction (*F*_(6,42)_ = 3.67, *p* < 0.005) and 3-way interaction (*F*_(6,42)_ = 3.05, *p* < 0.01). Block x infusion and infusion x lure interactions were not statistically significant (*F*_(2,14)_ = 1.39, *p* = 0.28 and *F*_(3,21)_ = 0.80, *p* = 0.51).

To further probe effects of muscimol after increased experience with mnemonic discrimination task procedures and stimuli, data for each test block were analyzed with separate repeated measures ANOVAs. The main effect of infusion condition was statistically significant in block 1 (*F*_(1,7)_ = 16.7, *p* < 0.005), though not in blocks 2 or 3 (*p*’s = 0.06 and 0.31, respectively). The interaction effect of infusion condition x lure, however, did not reach statistical significance for block 1 (*F*_(3,21)_ = 1.77, *p* = 0.18) indicating that the impairment following muscimol infusion when lure objects were novel was similar across all lure problems. The infusion × lure interaction did reach statistical significance for block 2 (*F*_(3,27)_ = 2.95, *p* < 0.05), and it is evident in Figure 5C that muscimol infusions during block 2 selectively impaired performance on the 70% target-lure overlap problem. During block 3, there was not a significant interaction effect of infusion condition and target-lure overlap (*F*_(3,27)_ = 0.78, *p* = 0.52). This was likely due to comparable performance following vehicle or muscimol infusion across all levels of target-lure overlap. As in prior analyses, significant main effects of lure were observed across all blocks (*p*’s < 0.001), such that performance decreased as feature overlap between target and lure increased.

### Exp.2: Muscimol impaired discrimination only when lures were novel

#### Discrimination performance

Given the finding that muscimol infusions in dorsal CA3/DG did not disrupt object discrimination performance by the final test block of Exp.1, we sought to determine if experience with lure objects on vehicle infusion days rendered the behavior resilient to disrupted hippocampal activity. Namely, did task performance become independent of dorsal hippocampus as animals accrued experience with the lures? To address this question, separate groups of rats received either a first block of 3 tests with hippocampal neural activity intact (veh-first, n=9), or with hippocampal neural activity disrupted (mus-first, n = 8; Fig.6A). Rats then received a second block of 3 tests with the reverse infusion condition (Fig.6A). Lure object stimuli were therefore comparatively novel in the first block of tests, and became increasingly familiar across the second block of tests. As for Exp.1, analyses revealed rats performed at chance on control trials with identical copies of the target object. A repeated measures ANOVA showed no difference in performance on these trials between veh-first and mus-first groups (*F*_(1,15)_ = 1.60, *p* = 0.23), across test blocks (*F*_(1,15)_ = 0.97, *p* = 0.34), and no group x block interaction (*F*_(1,15_) = 3.10, *p* = 0.10). One-sample t-tests also confirmed no difference from chance performance levels of 50% (*p*’s > 0.32). Data from these trials were thus excluded from subsequent analyses.

**Figure 6.**
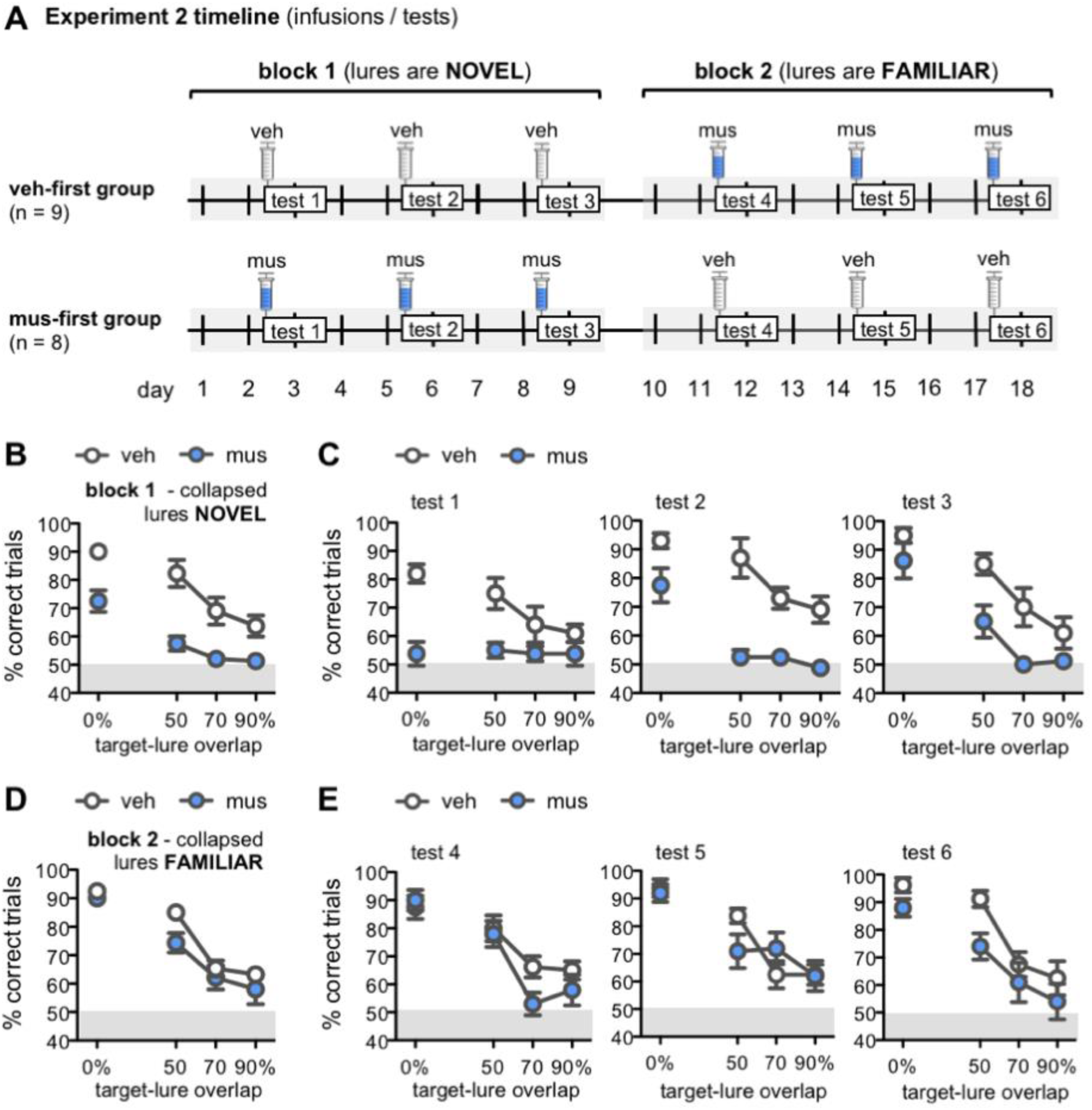
(**A**) Timeline of infusions and mnemonic discrimination tests administered in Experiment 2. Rats were randomly assigned after surgery to either receive vehicle infusions in the first test block (**veh-first group;** n=9) or muscimol infusions in the first test block (**mus-first group;** n=8). Each test block comprised 3 tests with the same infusion condition. In block 1, veh-first rats received vehicle and mus-first rats received muscimol prior to tests 1-3. In block 2, infusion conditions reversed; veh-first rats received muscimol and mus-first rats received vehicle prior to tests 4-6. (**B-E**) Performance (% correct trials) on mnemonic discrimination tests in Exp.2 plotted by trial type, with objects sharing 0, 50, 70, or 90% target-lure overlap. Test performance on days with vehicle infusions (veh) designated by open circles, and on days with muscimol infusions (mus) by filled circles. Data for block 1 are shown in (**B**) and (**C**). Data for block 2 are shown in (**D**) and (**E**). (**B/D**) Mean ± SEM performance on mnemonic discrimination tests collapsed across blocks 1 and 2, with a total of 30 trials per lure object for each infusion group. As in Exp.1, performance decreased as target-lure overlap increased (main effect lure: *F*_(3,45)_ = 61.8, *p* < 0.001). Muscimol impaired discrimination across all lures in block 1 (**B**), but did not impair discrimination in block 2 (**D**) (main effect block: *F*_(1,15)_ = 41.2, *p* < 0.001; block x group: *F*_(1,15)_ = 116.3, *p* < 0.001). (**C**) Mean ± SEM performance plotted for each test of block 1 (tests 1-3; 10 trials / lure object / infusion group). Muscimol infusions impaired discrimination across all tests of block 1, when lures were relatively novel (main effect group: *F*_(1,16)_ = 27.2, *p* < 0.001); however, rats in the mus-first group showed selective improvement across tests on trials with the more distinct lure objects (main effect test: *F*_(232)_ = 11.2, *p* < 0.001; test x lure: *F*_(6,96)_ = 6.13, *p* < 0.001; 3-way interaction: *F*_(6,96)_ = 2.86, *p* < 0.013). (**E**) Mean ± SEM performance plotted for each test of block 2 (tests 4-6; 10 trials / lure object / infusion group). Muscimol infusions did not impair discrimination across tests of block 2, when rats of the veh-first group had gained prior experience with lure objects (main effect lure: *F*_(3,45)_ = 58.5, *p* < 0.001 ; all other effects: *p*’s > 0.11).

Performance on mnemonic discrimination tests in Exp.2 is shown in Figure 6B-E. Data collapsed across tests of blocks 1 and 2 are shown in Figure 6B and 6D, respectively. A repeated measures ANOVA with infusion group (veh-first, mus-first) as a between subjects factor, and test block and lure object as within subjects factors showed a significant main effect of lure (*F*_(3,45)_ = 61.8, *p* < 0.001). This analysis also revealed a significant main effect of test block (*F*_(1,15)_ = 41.2, *p* < 0.001) and block x group interaction (*F*_(1,15)_ = 116.3, *p* < 0.001), but no significant main effect of group, block x lure interaction, or 3-way interaction (*p*’s > 0.08). Rats that received muscimol infusions in block 1 showed improved discrimination in block 2 when they received vehicle infusions (mus-first group; within subjects contrasts, α = 0.006; all *p*’s < 0.002). Conversely, rats that received vehicle in block 1 and muscimol in block 2 showed no change in performance between blocks (veh-first group; *p*’s > 0.05).

To clarify the improvement in discrimination performance observed with greater test experience and increasing familiarity of lure objects, data from individual tests of blocks 1 and 2 were compared with separate repeated measures ANOVAs (Fig.6C/E). Infusion group was entered as a between subjects factor, and test number (tests 1-3 for block 1, tests 4-6 for block 2), and lure object as within subjects factors. For tests of block 1 (Fig.6C), this analysis revealed significant main effects of lure object (*F*_(3,48)_ = 25.4, *p* < 0.001), test (*F*_(2,32)_ = 11.2, *p* < 0.001), and a test x lure interaction (*F*_(6,96)_ = 6.13, *p* < 0.001). Critically, discrimination performance differed between infusion groups on these first 3 tests (main effect of group: *F*_(1,16)_ = 27.2, *p* < 0.001; 3-way interaction: *F*_(6,96)_ = 2.86, *p* < 0.013). In contrast, for tests of block 2 (Fig.6E), only a significant main effect of lure was detected (*F*_(3,45)_ = 58.5, *p* < 0.001). All other effects and interaction terms did not reach statistical significance (*p*’s > 0.11).

In block 1, muscimol infusions impaired performance of mus-first rats relative to veh-first rats (Fig.6C). Specifically, on test 1, muscimol impaired performance on trials with 0% target-lure overlap (simple contrasts, α = 0.05/12 = 0.004; 0%: *F*_(1,16)_ = 29.1, *p* < 0.001; all other *p*’s > 0.008). On test 2, muscimol impaired discrimination on trials with each of the LEGO lure objects (50% overlap: *F*_(1,16)_ = 18.6, *p* < 0.001; 70% overlap: *F*_(1,16)_ = 22.0, *p* < 0.001; 90% overlap: *F*_(1,16)_ = 14.7, *p* < 0.001), though not the standard object (0% overlap: *F*_(1,16)_ = 6.67, *p* = 0.02). However, by test 3, no statistically significant differences were observed between infusion groups on any trial type (*p*’s > 0.008). In block 2, with infusion conditions reversed, muscimol did not impair performance in rats from the veh-first group (Fig.6E). No statistically significant differences in performance on trials with any lure were detected on test 4 (*p*’s > 0.04), test 5 (*p*’s > 0.09), or test 6 (Fig.6D; *p*’s > 0.01). Together, these data indicate the ability of intra-hippocampal muscimol infusions to impair mnemonic discrimination performance depends on relative novelty of the lure objects.

#### Response strategy and reaction latencies

Rats default to an egocentric side-biased response strategy (i.e. perseverative selection of the left- or right-presented object) during initial learning on object discrimination tasks (Lee and Solivan, 2008; Jo and Lee, 2010; Lee and Byeon, 2014; Hernandez et al., 2015, 2017; Johnson et al., 2017). While this effect is particularly pronounced in aged relative to young adult rats, it also emerges in young adult rats when tested with lure objects that share greater feature overlap with a previously learned target (Johnson et al., 2017). To determine if disruption of neural activity in dorsal CA3/DG influenced rats’ abilities to suppress this perseverative response strategy, side bias index values were compared across test sessions from Exp.2 (Fig.7A). This index is calculated as the absolute value of (total number of left well choices - total number of right well choices) / total number of trials completed. An index of 1 therefore reflects behavior governed entirely by a side bias, while an index of 0 reflects an equal number of responses made to each side. In block 1, mean side bias index values were greater in rats that received muscimol infusions, relative to those that received vehicle (main effect of infusion condition: *F*_(1,15)_ = 33.4, *p* < 0.001). However, side bias decreased across tests of this block irrespective of infusion condition, as rats gained experience with lure objects and test procedures (main effect of test: *F*_(2,30)_ = 6.38, *p* < 0.005; no infusion x test interaction: *F*_(2,30)_ = 2.41, *p* = 0.11). Conversely, in block 2, mean side bias index values did not differ based on infusion condition or test (*p*’s > 0.25). Within subjects contrasts showed side bias decreased from block 1 to 2 in rats that initially received muscimol infusions (*F*_(1,7)_ = 67.7, *p* < 0.001), but remained consistent across block 1 and 2 in rats that first received vehicle (*F*_(1,8)_ = 0.07, *p* = 0.80). Additional analyses confirmed greater side bias on trials with lures that shared greater feature overlap, as observed in prior studies (data not shown; main effects of lure: *p*’s < 0.001; Johnson et al., 2017). Nonetheless, side bias index values of rats that received muscimol in block 1 tests were greater across all trial types (tests 1-3, main effects of infusion condition: *p*’s < 0.04). This consequence of muscimol was not observed when rats received vehicle in block 1, gaining experience with task conditions and lure objects with hippocampal neural activity intact (tests 4-6, main effects of infusion condition: *p*’s > 0.11).

**Figure 7.**
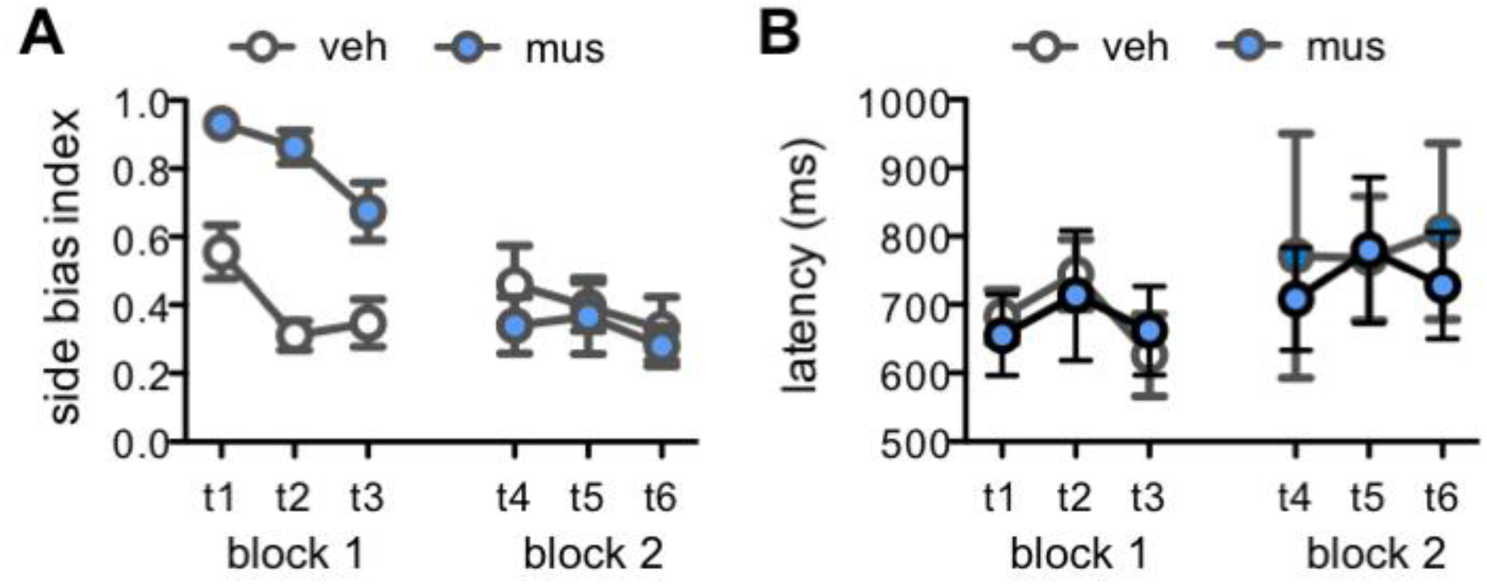
Measures of response selection behavior in Experiment 2. Data from days with vehicle infusions (veh) designated by open circles, and on days with muscimol infusions (mus) by filled circles. Data are plotted by test day (tests 1-6; t1-t6) across each of the two test blocks. (**A**) Mean ± SEM side bias index values across tests. A side bias index of 1 reflects a complete bias to select the object presented on either the left or right side of the choice platform, while a side bias of 0 reflects no side bias and an equal number of responses made to the left and right sides. In block 1, side bias index values were greater after muscimol relative to vehicle (main effect infusion: *F*_(1,15)_ = 33.4, *p* < 0.001), though decreased across tests as rats gained experience with lure objects and were increasingly able to suppress their inherent bias to correctly select the target object (main effect test: *F*_(2,30)_ = 6.38, *p* < 0.005). Conversely, in block 2, mean side bias index values did not differ based on infusion condition or test (*p*'s > 0.25). (**B**) Mean ± SEM response latency values for correct discrimination responses made to rats' inherently biased side. Latencies did not differ by infusion condition (main effect infusion: *F*_(1,9)_ = 0.61, *p* = 0.455) or across test days (main effect test: *F*_(5,45)_ = 0.81, *p* = 0.547; infusion x test: *F*_(5,45)_ = 0.21, p = 0.956).

To determine if muscimol infusions led to general behavioral slowing, or otherwise affected the duration of response selection, reaction latencies on correct trials were scored for each test session. Rats that received muscimol infusions in block 1 made few if any responses to their non-biased side, therefore data for correct responses to the non-biased side were not analyzed due to missing values. Mean reaction latencies for correct responses to the biased side were compared across test sessions (Fig.7B). These latencies to displace the target object when presented on the rats’ biased side did not differ by infusion condition (main effect of infusion: *F*_(1,9)_ = 0.61, *p* = 0.455) or across test days (main effect of test: *F*_(5,45)_ = 0.81, *p* = 0.547; infusion x test interaction: *F*_(5,45)_ = 0.21, *p* = 0.956). Together, these data suggest that muscimol infusions in block 1 led rats to default to their inherent side-biased response strategy when faced with the relatively challenging task of differentiating the known target from novel lure objects. However, accurate performance of muscimol-infused rats in block 2, that had prior experience on the task with hippocampal neural activity intact, corresponded with suppression of the inherent side bias. In addition, muscimol infusions did not lead to behavioral slowing when administered in either test block.

### Muscimol impaired spatial alternation performance after prior repeated infusions

One potential explanation for the behavioral results of Exps.1 and 2 is that a tolerance to the pharmacological actions of muscimol emerged after multiple infusions of the drug. Our analyses of *Arc* mRNA expression within the hippocampus subsequently confirmed this was not the case; excitation of DG was indeed noted in rats from Exps.1 and 2 infused with muscimol after they completed multiple rounds of behavioral testing and infusions (Fig.1C), and this same pattern of Arc expression was also observed in rats following a first and single infusion of muscimol relative to vehicle (Fig.3). Nevertheless, we sought to test the possibility that hippocampal muscimol infusions became less effective in modulating behavior over time in rats from Exp.2. Rats were trained on a spatial alternation task to a criterion of 10 consecutive correct alternations, then tested following a muscimol infusion, and tested once more following a vehicle infusion (Fig.8). Muscimol increased rats’ number of spatial errors within test day 1, relative to their pre-infusion baseline (paired samples t-test, *t*_(7)_ = −3.22, *p* < 0.01). In contrast, vehicle infusions did not alter rats’ alternation performance on test day 2 relative to pre-infusion baseline (*t*_(7)_ = −0.15, *p* = 0.86). Critically, comparisons confirmed performance did not significantly improve on pre-infusion baseline trials from test day 1 to day 2 (*t*_(7)_ = −0.52, *p* = 0.62), but was significantly impaired following muscimol infusion relative to vehicle infusion (*t*_(7)_ = −2.60, *p* < 0.04).

**Figure 8.**
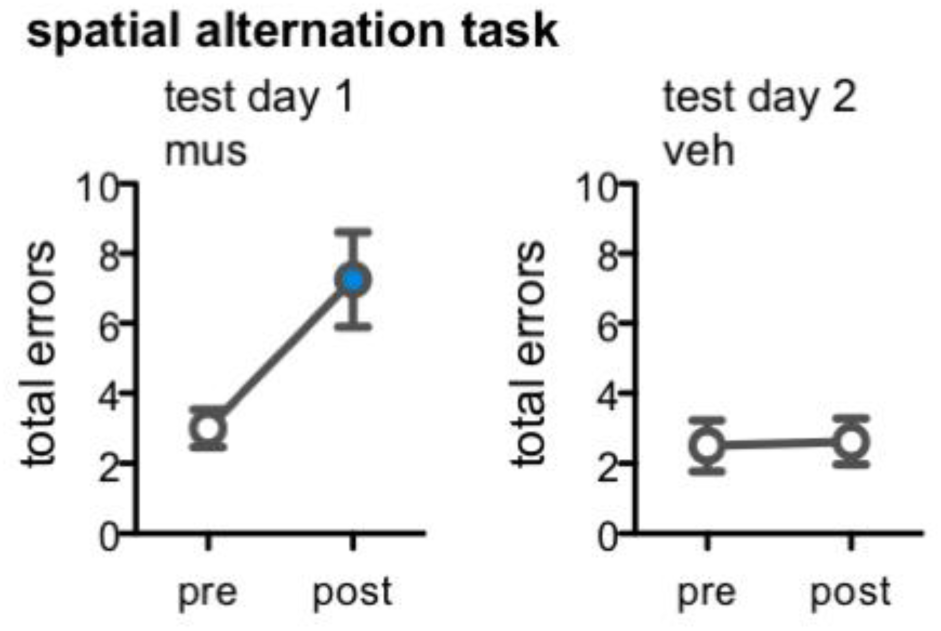
Effect of dorsal CA3/DG muscimol infusions on spatial alternation performance, as a behavioral control condition in rats from Exp. 2 (n=8). Plots show the total number of spatial memory errors (i.e., return to same arm of the figure-8 maze on 2 consecutive laps, rather than alternating between the two arms) on 10 pre-training trials and 10 post-infusion trials on test day 1 (all rats infused with muscimol; mus) and test day 2 (all rats infused with vehicle; veh). Mean ± SEM number of errors for pre-training and post-vehicle trials designated by open circles, and for post-muscimol trials with filled circle. Muscimol impaired alternation performance, even after completing all infusions and mnemonic discrimination tests of Exp.2 (*t*_(7)_ = −3.22, *p* < 0.01); vehicle did not alter rats’ alternation performance (*t*<_(7)_ = −0.15, *p* = 0.86).

## DISCUSSION

The current studies were designed to test the role of CA3/DG in object discrimination in young adult rats. In particular, based on human neuroimaging data linking CA3/DG activity to accuracy in distinguishing highly similar visual stimuli (Kirwan and Stark, 2007; Bakker et al., 2008; Yassa and Stark, 2011; Reagh and Yassa, 2014; Reagh et al., 2018), we sought to determine if disrupting neural activity in the rodent CA3/DG with intrahippocampal infusions of muscimol would selectively impair discrimination of a target object from highly similar lure objects. However, in verifying the effects of muscimol on neural activity, we discovered infusion of this GABAA agonist at the coordinates used here produced excitation throughout the DG granule cell layer (Fig.1C), which typically shows sparse levels of activation assessed with the same methods (Small et al., 2004; Marrone et al., 2011, 2012; Penner et al., 2011; Satvat et al., 2011; Gheidi et al., 2013). A subsequent experiment confirmed that, relative to vehicle, muscimol infusions robustly *increased* expression of the activity-dependent immediate-early genes *Arc* and *Homer1a* in DG (Fig.3). Although we have previously found that muscimol infusions in the perirhinal and prefrontal cortices reduced *Arc* expression (Hernandez et al., 2017), surprisingly this was not the pattern observed here. A previous study that infused muscimol into the hippocampus at different coordinates than those used here did report a reduction in *Arc* expression in CA1 (Kubik et al., 2012). The apparent discrepancy between this earlier result and the current experiments could be explained by the position of infusions, both more anterior and lateral to the DG (Kubik et al., 2012). The hyperexcitation of DG granule cells observed here most likely arose through muscimol’s actions on the dense populations of interneurons present in the hilus and DG molecular layers, which when inactivated by muscimol released the DG granule cell population from what is normally strong inhibition (Amaral and Lavenex, 2007; Amaral et al., 2007; Treves et al., 2008). Though it is difficult to say whether previous studies using the same approach may have unintentionally brought about similar patterns of DG excitation, our results underscore the importance of verifying both the behavioral consequences of drug infusions as well as the effects on neural activity.

Although muscimol infusions led to hyperexcitation in dorsal DG granule cells, the results of Exps.1 and 2 revealed a clear effect of disrupting hippocampal neural activity in this manner on discrimination performance. Theory and computational models have proposed that one critical role of the DG is to support orthogonalization of inputs from parahippocampal cortical regions, minimizing interference (McNaughton and Morris, 1987; Treves and Rolls, 1992; O’Reilly and McClelland, 1994; Rolls and Kesner, 2006; Treves et al., 2008; Yassa and Stark, 2011; Santoro, 2013; Kesner and Rolls, 2015; Rolls, 2016). This function is historically attributed to relatively sparse activity in DG, through tight inhibitory control from adjacent interneuron populations, and to anatomical organization of projections from the entorhinal cortex, where few afferent neurons project onto many granule cells (Witter et al., 1989; Amaral and Lavenex, 2007; Amaral et al., 2007; Treves et al., 2008). Based on these assumptions, it is plausible that increasing excitability of DG granule cells would have an equivalent if not greater impact on rats’ abilities to distinguish similar stimuli than silencing DG. This prediction was in part borne out, as hyperexcitability of granule cells impaired target-lure discrimination performance across *all levels* of target-lure overlap (including the 0% overlap problem). Moreover, impairments were most profound when lures were novel, and increasing experience with target-lure discrimination problems led to behavioral improvements even when DG activity was disrupted. These observations suggest that, contrary to prevailing models of DG function, computational contributions of DG granule cells extend beyond the orthogonalization of similar inputs and may be modulated by stimulus novelty. In fact, recent neurophysiology data provide evidence questioning the role of granule cells for performing pattern separation computations. Specifically, in optogenetically-verified neuron populations, DG granule cells have been shown to not exhibit updated activity patterns when the testing arena in a common room is changed (Senzai and Buzsáki, 2017). These data suggest DG granule cells may not support the disambiguation of overlapping sensory input, but instead help discriminate novel stimuli from those that are familiar across all levels of sensory overlap. A potential role for the DG in discriminating between novel and familiar stimuli may not be surprising considering the strong input this region receives from the locus coeruleus (Blackstad et al., 1967; Haring and Davis, 1985; Amaral and Witter, 1995). Locus coeruleus neuron firing is modulated by novelty (Vankov et al., 1995), and noradrenaline modulates the excitability (Kitchigina et al., 1997) and plasticity (Harley, 1991) of DG granule cells. Under this framework, when a lure is novel, functional connectivity between the locus coeruleus and DG would be critical for discrimination. As the lure is learned, however, other networks can support the behavior independent of DG.

Another potential explanation for the current data are that disruption of hippocampal neural activity impeded general behavioral performance or procedural knowledge of the task. This was particularly concerning given the possibility that rats may have been experiencing sub-convulsive seizures following infusions with muscimol. Importantly, during behavioral experiments, in which performance was video recorded and scored offline, no rats displayed overt seizures and all rats performed normally on task procedures. Moreover, analyses of reaction times showed no effect of hippocampal muscimol on general activity or sensorimotor abilities. After muscimol infusions rats were in fact quicker to select responses and showed less deliberation between objects. Calculation of side bias index across test days revealed that, when infused with muscimol, rats defaulted to a side-biased strategy. This recapitulates the suboptimal response strategy employed by aged rats when distinguishing lures that share greater feature overlap (Johnson et al., 2017), and by young adult rats when hippocampal or medial temporal lobe activity is disrupted in similar forced-choice object discrimination tasks (Lee and Solivan, 2008; Jo and Lee, 2010; Hernandez et al., 2017). Taken together, these data confirm muscimol and its actions in the DG did not result in gross behavioral impairment, but rather led to selective deficits in discriminating a target object from novel lures.

Findings of the current experiments differ from those of previous studies using rodent models to investigate the contributions of DG to stimulus discrimination. One key difference is in methodology; prior experiments have disrupted hippocampal activity by lesioning DG granule cells with the neurotoxin colchicine (Gilbert et al., 2001; Morris et al., 2012; Weeden et al., 2014; Oomen et al., 2015). Colchicine lesions produce permanent neuronal damage, precluding the possibility of gaining experience in behavioral tasks post-op with DG activity intact. Prior investigations adopting this approach have shown colchicine lesions of the dorsal DG cause sustained impairments in tasks that involve discrimination of spatial locations on an open platform maze (Gilbert et al., 2001) or a two-dimensional touchscreen (Oomen et al., 2015). Gilbert and colleagues (2001) found that rats previously trained to resolve target locations from lure locations were impaired in distinguishing locations separated by 60 cm or less when re-tested after DG lesions (Gilbert et al., 2001). Rats were not impaired in distinguishing locations with greater spatial separation, 82.5-105 cm apart (Gilbert et al., 2001). It is interesting to note that a subsequent study using identical apparatus and procedures showed ibotenic acid lesions of dorsal CA3 impaired discrimination of locations across all spatial separations (Gilbert and Kesner, 2006). Given these data, it is conceivable that DG granule cell hyperactivity disrupted neural firing in CA3, without altering *Arc* expression, and combined CA3/DG dysfunction following muscimol led to the observed behavioral impairment.

Using an automated touchscreen non-match to location task, Oomen et al. (2015) found DG lesions produced greater impairments on a version that posed a greater working memory burden, with variable intra-trial delay within sessions or requiring selection of the target location from 3 versus 2 choices (Oomen et al., 2015). Moreover, DG lesions slowed acquisition of the 3-choice version of the task, and impaired performance on probe tests in which the distance between target (S+) and lure (S-) locations was manipulated. Similar to the current studies, target-lure discrimination across all distances was impaired following DG lesions (Oomen et al., 2015). One other study has investigated the effect of dorsal DG lesions on spatial discrimination, though without pre-surgical training (Morris et al., 2012). Morris and colleagues (2012) found DG lesions impaired acquisition of reward-place associations on a radial arm task when reward and non-reward arms were adjacent to each other, but not when separated by multiple arms. In addition, employing stimuli from a different sensory modality, Weeden and colleagues (Weeden et al., 2014) found ventral DG lesions impaired olfactory discrimination, though only for similar odorants and at longer intra-trial delays. A pronounced difference of our results from prior studies is that disrupting DG activity with muscimol infusions *did not impair* discrimination performance in rats that gained prior experience with task stimuli. This finding suggests that as the lures become familiar, neural activity across a broader cortico-hippocampal network is able to support task performance when DG function is disrupted, possibly through direct projections from entorhinal, perirhinal, and postrhinal cortex to CA3 and CA1 (Burwell et al., 1995).

Neuroimaging data have shown that increased activity in CA3/DG in aged relative to young subjects correlates with impaired mnemonic discrimination performance (Yassa et al., 2011a, 2011b; Reagh et al., 2018). While it may appear in the current studies that muscimol infusions recapitulated this effect of age, hippocampal hyperactivity observed in older adults is largely believed to arise from CA3. For instance, though fMRI does not afford the spatial resolution to isolate BOLD signals from DG versus CA3, invasive recordings in rats (Wilson et al., 2005, 2006; Robitsek et al., 2015) and monkeys (Thomé et al., 2016), slice recordings in rats (Simkin et al., 2015), and cellular imaging in rats (Maurer et al., 2017) have shown CA3 pyramidal cells are hyperactive in aged animals. This is consistent with age-associated dysfunction within hilar interneurons (Spiegel et al., 2013; Koh et al., 2014). In contrast, DG is less active in aged humans, monkeys and rats (Small et al., 2002, 2004, 2011). In the current data, *Arc* and *Homer1a* levels in CA3 were comparable following vehicle and muscimol infusions, and it was only DG granule cells that exhibited hyperactivity. Thus, the muscimol infusions used here produced a pattern of neural activity that was more consistent with sub-convulsive temporal lobe seizures, which are associated with granule cell discharges (Dengler et al., 2017; Bumanglag and Sloviter, 2018), and observed in individuals with Alzheimer’s disease (Palop and Mucke, 2009, 2010; Vossel et al., 2013, 2016). Although repeated granule cell discharges and resultant epileptogenesis could potentially produce hippocampal sclerosis, findings that performance improved over repeated tests and that CA3 activity was not elevated suggest that acute granule cell hyperactivity induced by muscimol did not propagate or elicit a chronic imbalance between excitation and inhibition.

In conclusion, the current studies emphasize the importance of verifying effects of experimental manipulations with both behavioral and neural read-outs. By assessing the expression of activity-dependent immediate-early gene transcripts after muscimol infusions for routine histology, we found these infusions targeting dorsal CA3/DG led to widespread hyperexcitation of the DG granule cell layer. Nonetheless, disrupting DG activity in young adult rats with this approach impaired discrimination of a target object from novel lure objects in a rodent version of the mnemonic similarity task (Stark et al., 2015; Huffman and Stark, 2017; Johnson et al., 2017; Stark and Stark, 2017). Repeated testing across multiple infusion blocks showed muscimol profoundly impaired mnemonic discrimination when object stimuli were novel, but no longer impaired discrimination when stimuli became familiar despite continued disruption of DG function. These findings suggest intact hippocampal function is most critical to mnemonic discrimination when faced with new information, while, with learning, other networks are capable of supporting these abilities independent of DG (Stern et al., 2001). Recent fMRI data indicate activity in parahippocampal, retrosplenial, and occipito-temporal cortical regions can be linked to accurate mnemonic discrimination (O’Neil et al., 2009; Ryan et al., 2012; Reagh and Yassa, 2014; Bakker et al., 2015; Pidgeon and Morcom, 2016; Reagh et al., 2017, 2018). Furthermore, reduced integrity of the hippocampal cingulum bundle, which connects medial temporal and frontal cortical regions, is associated with impaired mnemonic similarity task performance in older adults (Bennett and Stark, 2016). To our knowledge, frontal cortical contributions to similarity-dependent object discrimination have not yet been assessed in animal models. It will be of interest to delineate in future studies how relative contributions of medial temporal, frontal, and hippocampal regions shift with increasing experience.

## Acknowledgements

This work was supported by the McKnight Brain Research Foundation, National Institute on Aging grants K99AG058786 (SAJ), R01AG049722 (SNB), R03AG049411 (SNB, APM, JLB), McKnight Brain Institute Fellowship (SAJ), UF Research Seed Opportunity Fund, and UF University Scholars Program Award (SMT). We thank Sai Balusu, Katelyn Carty, Shiela Chancoco, Deandra Chetram, Gabriela Colon, Nardin Derias, Courtney Desrosiers, Najla Faddoul, Chelsea Feuer, Amanda Gramacy, Stephanie Guadarrama, Juan Leon, Hyeran Grace Oh, Maria Pava, Jordan Reasor, Lindsay Santacroce, Leila Shafiq, and Sabrina Zequeira for their help with carrying out behavioral experiments and analyses of video recordings. We also thank Abbi Hernandez for assistance in collecting tissue for histology.

## Notes

The authors declare no conflict of interest.

